# Transit and integration of extracellular mitochondria in human heart cells

**DOI:** 10.1101/157164

**Authors:** Douglas B. Cowan, Rouan Yao, Jerusha K. Thedsanamoorthy, David Zurakowski, Pedro J. del Nido, James D. McCully

**Affiliations:** Department of Anesthesiology, Perioperative and Pain Medicine, Boston Children’s Hospital, Boston, Massachusetts 02115, USA.; Department of Anæsthesia, Harvard Medical School, Boston, Massachusetts 02115, USA.; Harvard Stem Cell Institute, Cambridge, Massachusetts 02138, USA.; Department of Cardiac Surgery, Boston Children’s Hospital, Boston, Massachusetts 02115, USA.; Department of Surgery, Harvard Medical School, Boston, Massachusetts 02115, USA.

**Author notes:** Douglas B. Cowan, Boston Children’s Hospital, 300 Longwood Avenue, Enders 312.1, Boston, MA 02115-5724 USA., E-mail –.

## Abstract

Tissue ischemia adversely affects the function of mitochondria, which results in impairment of oxidative phosphorylation and compromised recovery of the affected organ. The impact of ischemia on mitochondrial function has been most extensively studied in the heart because of the morbidity and mortality associated with injury to this organ. Because conventional methods to preserve cell viability and function following an ischemic injury are limited in their efficacy, we developed a unique approach to protect the heart by transplanting respiration-competent mitochondria isolated from a non-ischemic tissue to the ischemic region. Our experiments in animals have shown that transplantation of isolated mitochondria to injured heart tissue leads to decreases in cell death, increases in energy production, and improvements in contractile function. We also discovered that exogenously-derived mitochondria injected or perfused into ischemic hearts were readily internalized by cardiac cells through actin-dependent endocytosis. Here, we describe the use of three-dimensional super-resolution microscopy and transmission electron microscopy to determine the intracellular fate of exogenous mitochondria in non-dividing human iPS-derived cardiomyocytes and dividing primary human cardiac fibroblasts. We show isolated mitochondria are internalised in human cardiac cells within minutes and then transported to endosomes and lysosomes. The majority of exogenous mitochondria escape from these compartments and fuse with the endogenous mitochondrial network, while some organelles are degraded through hydrolysis. Understanding this process may guide the development of treatments directed at replacing or augmenting impaired mitochondria in ischemic tissues and provide new options to rejuvenate dysfunctional mitochondria in a wide range of human diseases and disorders.

## Introduction

Mitochondria play an essential role in energy production and cellular homeostasis. Dysfunction of these organelles as a result of ischemia or genetic mutations can lead to the loss of high-energy phosphate reserves, accumulation of mitochondrial calcium, and a buildup of reactive oxygen molecules^1–5^. Our previous studies demonstrated that transplanting isolated mitochondria to the ischemic heart leads to reductions in infarct size, increases in adenosine triphosphate (ATP) production, and improvements in contractility^6,7^. We also observed that mitochondria injected or perfused into the heart were rapidly internalised by a variety of cardiac cells including cardiomyocytes and fibroblasts^7,8^. Additional experiments using cell cultures proved that the uptake of mitochondria occurs through actin-dependent endocytosis and results in rescue of cellular function by increasing energy production and replenishing mitochondrial DNA (mtDNA)^9^. While other researchers have also observed endocytic uptake of extracellular mitochondria, the exact intracellular trafficking and fate of these organelles remains unknown^10–15^.

In this study, we used three-dimensional super-resolution structured illumination microscopy (3-D SR-SIM) and transmission electron microscopy (TEM) to reveal the intracellular fate of exogenous mitochondria in human induced pluripotent stem cell-derived cardiomyocytes (iPS-CMs) and human cardiac fibroblasts (HCFs). By labelling extracellular mitochondria with fluorescent proteins or gold nanoparticles, we were able to observe the transit of endocytosed mitochondria in these cells. Distinct fluorescent labelling of various cell compartments in iPS-CMs and HCFs allowed us to visualise the progression of exogenous mitochondria through the endolysosomal system and established that these organelles primarily integrate with the endogenous mitochondrial network in both cell types. When combined with the findings of other investigators, our results strongly support the notion that the uptake and subsequent fusion of extracellular mitochondria with recipient cell mitochondria is an evolutionarily-conserved and pervasive biological process^7–16^. A thorough understanding of the intracellular fate of exogenous mitochondria may present new treatment strategies for the ischemic heart and drive the development of organelle-based therapeutics for a host of other human diseases and disorders^17–20^.

## Results

### Labelling of organelles and characterization of isolated mitochondria

We investigated the temporal and spatial fate of endocytosed mitochondria in non-dividing iPS-CMs and dividing HCFs. The identity and morphology of these cardiac cells was substantiated by immunostaining with α-actinin (ACTN) and vimentin and both cell types were shown to react well with established mitochondrial antibodies (TOMM20 or MTC02) (Extended Data Fig. 1a). To discern exogenous mitochondria within cultured cells, we labelled HCF mitochondria with green fluorescent protein (GFP) and used red fluorescent proteins (RFP) to label various HCF and iPS-CM cell compartments through baculovirus-mediated transfer of mammalian fusion genes (Fig. 1a). Both cell types were readily infected with baculoviruses carrying fluorescent protein genes and demonstrated specific expression of GFP or RFP in organelles including mitochondria, early and late endosomes, lysosomes, Golgi complexes, and the endoplasmic reticulum (Extended Data Fig. 1b). Isolated HCF GFP-labelled mitochondria were also stained with MitoTracker Red CMXRos or a human mitochondria-specific antibody (MTC02) to confirm their identity and then imaged using 3-D SR-SIM (Fig. 1b).

**Figure 1.**
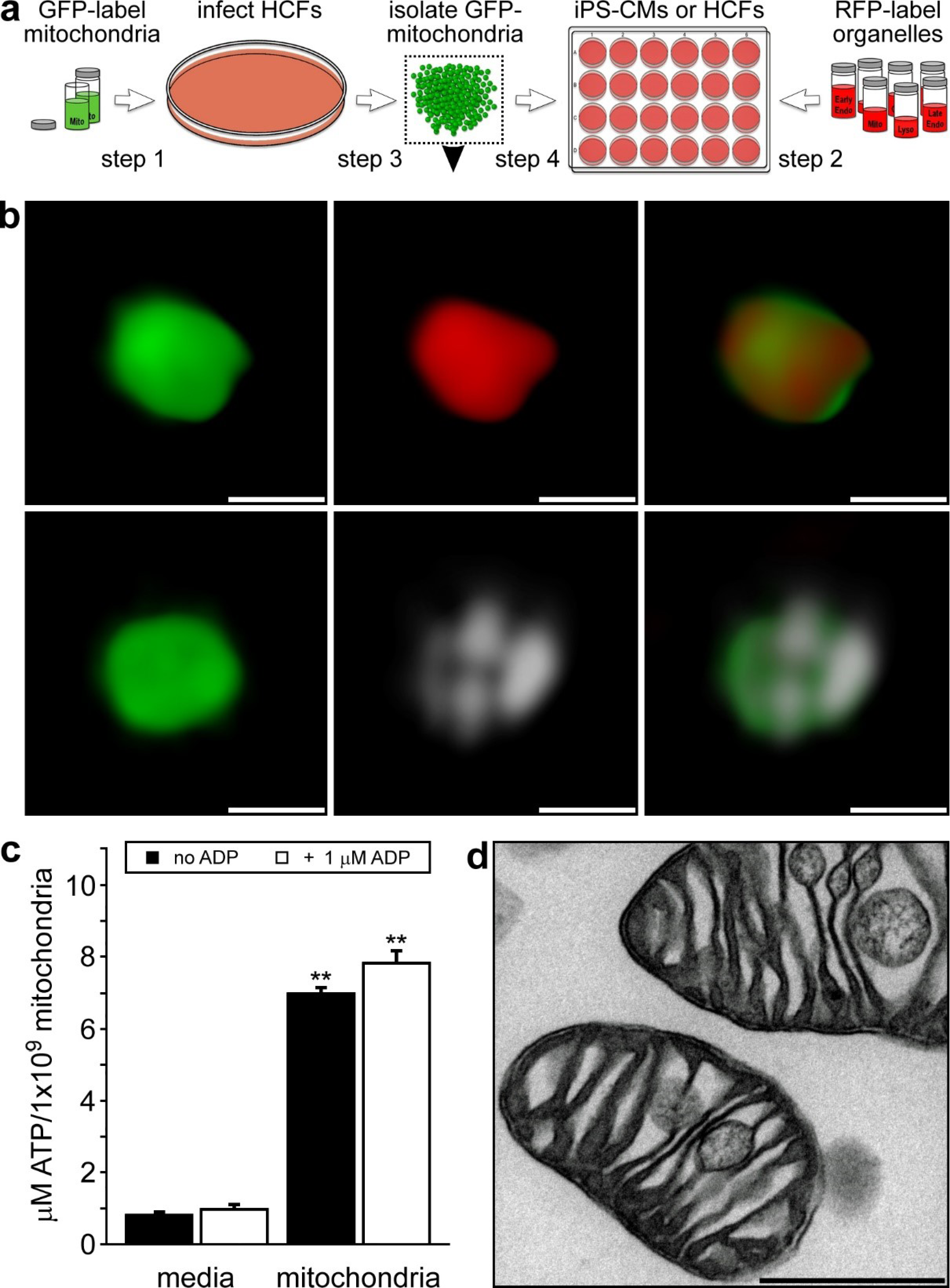
Experimental strategy and characterisation of isolated human fibroblast mitochondria. **a**, HCFs infected with BacMam CellLight Mitochondria-GFP were used for mitochondrial isolations and iPS-CMs or HCFs on coverslips were infected with RFP CellLight reagents to label specific cell structures. Isolated GFP-labelled mitochondria were added to RFP-labelled cells for 0 to 4 h. **b**, 3-D SR-SIM of isolated HCF mitochondria stained with MitoTracker Red CMXRos (top panels) or the human mitochondria-specific antibody MTC02 (bottom panels). Isolated mitochondria, stained mitochondria, and the combined image are shown left to right. Scale bars, 0.5 μm. **c**, ATP content in media or isolated HCF mitochondria in the presence or absence of 1 μM ADP. Error bars indicate standard error of the mean. ATP levels were significantly higher in mitochondria than in media-only control groups (Student’s t-test, ** p < 0.001). **d**, transmission electron microscopy of isolated HCF mitochondria. Scale bar, 0.5 μm. The image is representative of four separate mitochondrial isolations.

Isolated mitochondria were generally spherical in shape and varied in diameter from 250 to 2000 μm with the majority of these organelles falling within the 350 to 600 μm range^7^. In addition, adenosine triphosphate (ATP) measurements verified that isolated mitochondria were viable and energised (Fig. 1c), whereas transmission electron microscopy revealed these organelles were structurally intact (Fig. 1d)^7,8,21^.

### Endocytosis and intracellular position of exogenous mitochondria

To examine endocytosis of isolated mitochondria in iPS-CMs, we first labeled HCF mitochondria with 10 nm diameter gold nanoparticles (Fig. 2a) to allow for their identification by transmission electron microscopy. Labelled exogenous mitochondria were observed outside cells, near cell surfaces, in the process of endocytic engulfment, and inside iPS-CMs^22^. Next, the intracellular position of endocytosed mitochondria in iPS-CMs was determined using four color 3-D SR-SIM (Fig. 2 and Extended Data Fig. 2). Because we did not observe GFP-labelled mitochondria associated with either the Golgi complex or endoplasmic reticulum, we focused on examining whether RFP-labelled endosomes, lysosomes, or endogenous mitochondria colocalised with internalised HCF mitochondria. Early endosomes (Fig. 2b), late endosomes (Fig. 2c), and lysosomes (Fig. 2d) of cardiomyocytes were all found to contain GFP-labelled mitochondria at each time examined (0.5, 1, 2, and 4h). Endocytosed mitochondria were also found in close proximity to the endogenous mitochondrial network, which was generally located apical to the contractile apparatus (Extended Data Fig. 2a).

**Figure 2.**
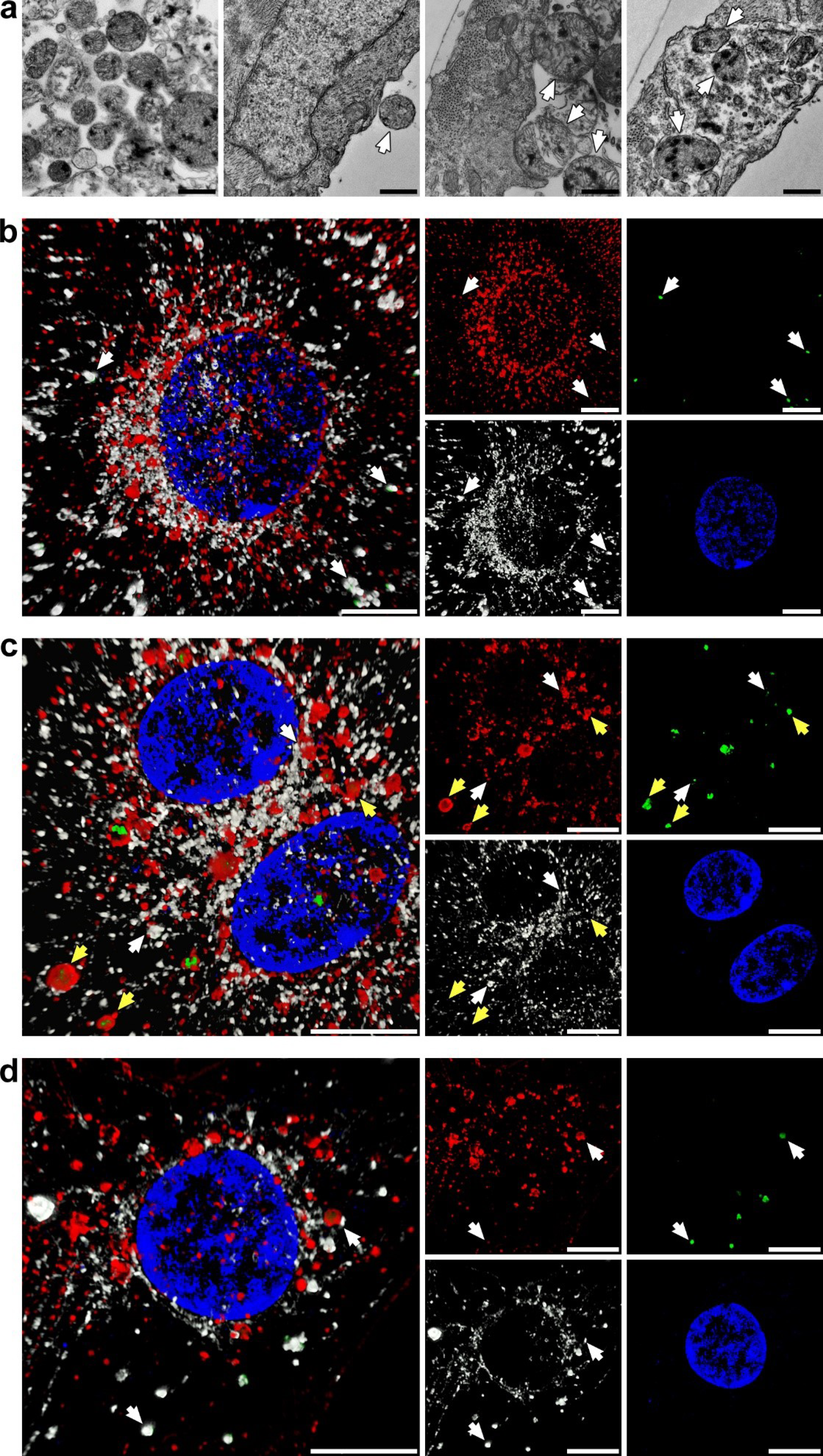
Uptake and endocytic transport of extracellular mitochondria in cardiomyocytes. **a**, iPS-CMs were exposed to gold-labelled HCF mitochondria for 2.5, 5, and 10 min and imaged by TEM. Labelled mitochondria had electron dense deposits and were apparent at all three study times outside cells, adjacent to the apical cell surface, undergoing endocytosis, and inside cells (left to right and indicated by arrows). Images are representative of four experiments at 5 min. Scale bars, 0.5 μm. **b**, Four color 3-D SR-SIM of colocalisation of exogenous mitochondria (green) with early endosomes (red) at 4 h. Nuclei were stained with DAPI (blue) and both endogenous and exogenous mitochondria were detected with the MTC02 antibody (white). The combined image (left) and each color (right) are shown. Scale bars, 10 μm. **c**, Colocalisation of HCF mitochondria (green) with late endosomes (red) at 1 h. Scale bars, 10 μm. **d**, Colocalisation of mitochondria (green) with lysosomes (red) at 4 h. Scale bars, 10 μm. In **c-d**, white arrows indicate examples of exogenous mitochondria associated with each compartment stained with the anti-mitochondria antibody and yellow arrows show internalised, GFP-labelled mitochondria encapsulated within late endosomes that were not stained with MTC02. Nuclei were stained with DAPI (blue). Images represent twelve separate volumes analysed at each time (0.5, 1, 2, and 4 h).

### Escape of exogenous mitochondria from the endolysosomal system

Exogenous mitochondria associated with early endosomes, late endosomes, and lysosomes were observed to be fully encapsulated or partly contained within these compartments. Mitochondria partially enclosed in endocytic vesicles were considered to be escaping from their respective endosomal or lysosomal compartments. Cell preparations were also stained with anti-human mitochondria antibody (MTC02) to confirm that green fluorescence was attributable to the presence of intact organelles; though, this antibody was less reactive with encapsulated mitochondria — likely a consequence of reduced antigen accessibility. To further corroborate the identity of the above compartments, we stained cardiomyocyte cell preparations with early endosome antigen 1 (EEA1) and the late endocytic marker Rab7 (Extended Data Fig. 2b-d). In addition, we observed some lysosomes associated with exogenous mitochondria also stained for Rab7, which could simply epitomize the normal heterogeneity of the lysosomal compartment or indicate these particular compartments may be more accurately described as endolysosomal vesicles^23,24^.

The escape of exogenous mitochondria from endosomes and lysosomes was assessed by counting colocalisation of GFP-labelled mitochondria with each of these compartments. High-magnification 3-D SR-SIM indicated GFP-labelled mitochondria became increasingly reactive with MTC02 antibody upon emerging from late endosomes (Fig. 3a) as well as early endosomes and lysosomes (not shown). Enumeration of colocalisation of endocytosed mitochondria with endosomes and lysosomes at each time established that a majority of these exogenous organelles were associated with these vesicles at the earlier times (0.5 to 2 h); however, there was no obvious pattern to endolysosomal trafficking other than a decline in colocalisation at 4 h (Fig. 3b). The number of mitochondria escaping from these compartments showed the opposite trend with the largest number of organelles emerging from late endosomes and lysosomes at 4 h (Fig. 3c). We extended these analyses to include colocalisation between endogenous and exogenous mitochondria (Fig. 3d). These experiments demonstrated a large percentage of GFP-labelled mitochondria were closely associated with the cardiomyocyte mitochondrial network from 0.5 to 4 h. The functional consequence of the uptake and trafficking of extracellular mitochondria in cardiomyocytes is a sustained and significant increase in ATP production, similar to our earlier findings^9^. Together, these results indicate that endocytosis and intracellular trafficking of extracellular mitochondria in cardiomyocytes is an ongoing and efficient biological process.

**Figure 3.**
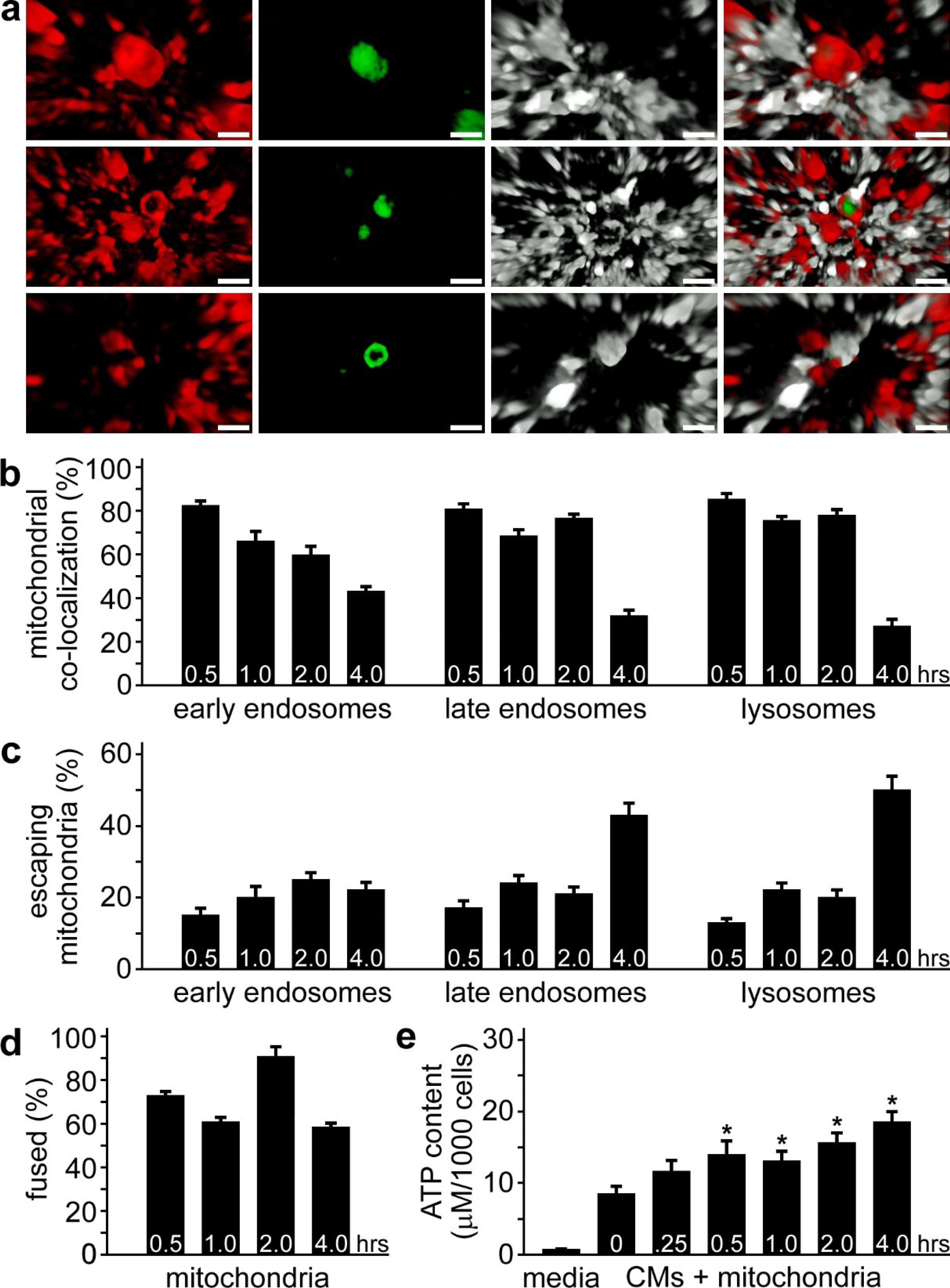
Encapsulation and escape of internalised mitochondria from cell compartments. **a**, Late endosomes (red) associated with GFP-labelled exogenous mitochondria (green) were imaged by 3-D SR-SIM. Examples of encapsulated and escaping mitochondria are shown after 1 h of incubation of isolated mitochondria and iPS-CMs. Images are representative of four experiments. Scale bars, 0.5 μm. **b**, quantitation of mitochondrial colocalisation with early endosomes, late endosomes, and lysosomes at 0.5, 1, 2, and 4 h. Results are expressed as the percentage of mitochondria associated with each compartment ***(i.e.*** encapsulated and escaping). **c**, quantitation of mitochondria escaping early endosomes, late endosomes, and lysosomes at 0.5, 1, 2, and 4 h. Results are expressed as the percentage of mitochondria associated with each compartment, but not fully encapsulated and represent a minimum of twelve experiments. **d**, quantitation of the colocalisation of endocytosed mitochondria with endogenous cardiomyocyte mitochondria. Results are expressed as the percentage of GFP-labelled mitochondria associated with RFP-labelled mitochondria. **e**, ATP content of culture media and cardiomyocytes treated with isolated HCF mitochondria for 0, 0.25, 0.5, 1, 2, and 4 h. ATP content is expressed as μM per 1000 cells and error bars indicate standard error of the mean. ATP was significantly higher in iPS-CMs treated with mitochondria for 0.5, 1, 2, and 4 h compared to untreated cells (Student’s t-test, * p < 0.02). Results were obtained from twelve separate experiments.

### Fusion of exogenous mitochondria with the endogenous mitochondrial network

The fusion of endocytosed mitochondria with the endogenous mitochondria of cardiomyocytes (Fig. 4a) and cardiac fibroblasts (Extended Data Fig. 3a-c) was assessed by 3-D SR-SIM. In combination with depth coding analyses (Extended Data Fig. 4a), these experiments established that exogenous mitochondria fuse with the endogenous mitochondrial network after escape from endolysosomal compartments and that mitochondrial fusion was evident at all of the studied times (0.5, 1, 2, and 4 h). The involvement of fusion proteins in facilitating this process was determined by studying expression of mitofusin-1 (MFN1), mitofusin-2 (MFN2), and optic atrophy 1 (OPA1) proteins in iPS-CMs and HCFs by immunoblot analysis (Fig. 4b). We then examined the spatial distribution of mitofusin proteins using 3-D SR-SIM (Fig. 4c and Extended Data Fig 4b). Mitofusin proteins are involved in tethering and fusing the mitochondrial outer membranes and OPA1 functions, in part, to enable inner mitochondrial membrane fusion^25–27^. The anti-mitochondria antibodies 113-1 (shown) or MTC02 were used to reflect the total mitochondrial content in iPS-CMs and HCFs^8^.

**Figure 4.**
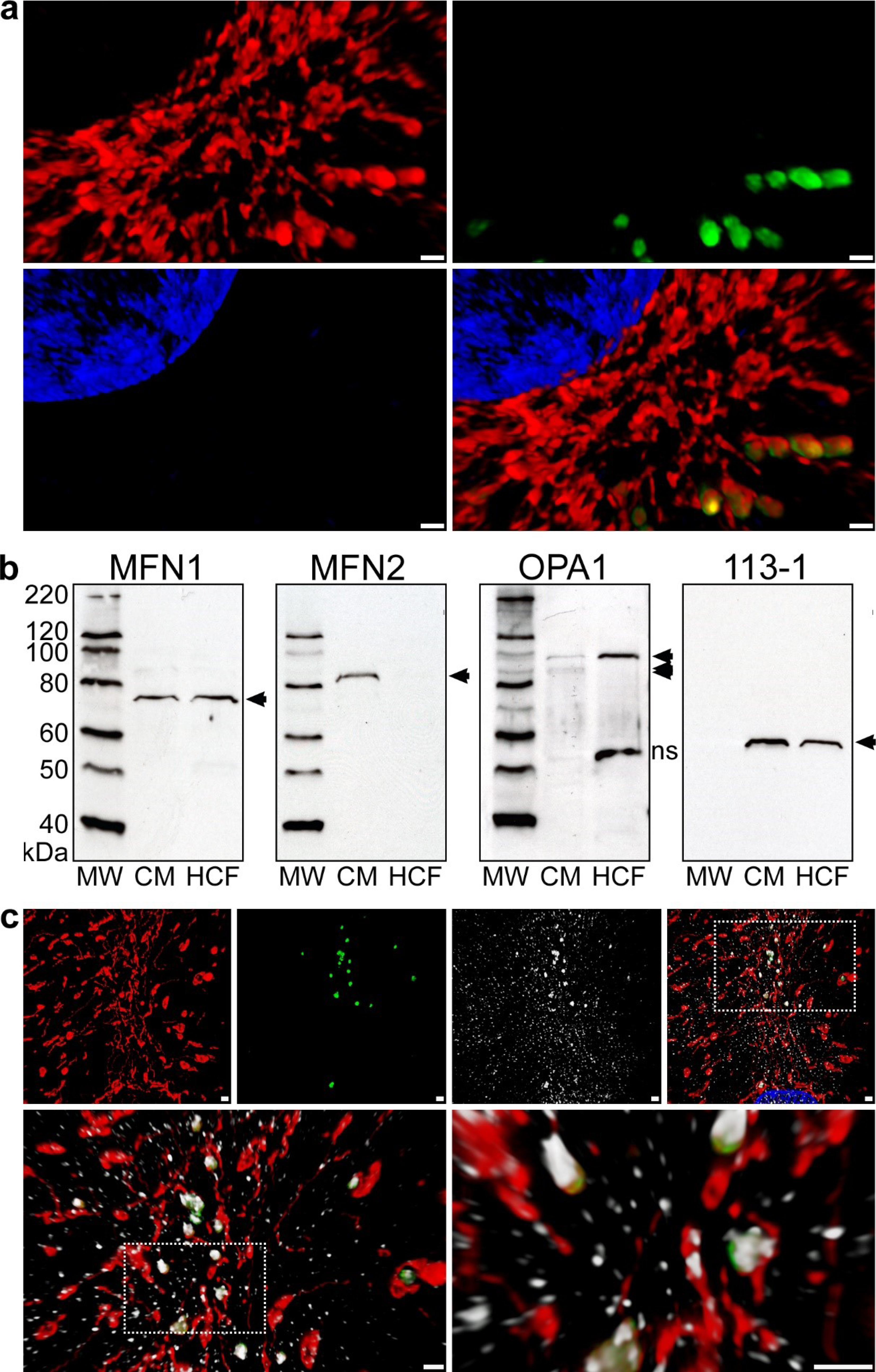
Fusion of exogenous mitochondria with the cardiomyocyte mitochondrial network. **a**, iPS-CMs with RFP-labelled mitochondria (red) treated for 4 h with isolated GFP-labelled mitochondria (green). After fixation, the nuclei were stained with DAPI (blue) and slides were imaged using 3-D SR-SIM. Individual red, green, and blue channels are shown along with the combined color rendering. Fusion of endocytosed mitochondria with the endogenous mitochondrial network is readily apparent. Mitochondrial fusion was apparent at all times examined (0.5, 1, 2, and 4 h) and four separate experiments were performed. Scale bars, 0.5 μm. **b**, immunoblot analysis of cell lysates (25 μg per lane) from iPS-CMs and HCFs using antibodies directed against MFN1, MFN2, OPA1, and mitochondria (113-1). Lanes loaded with molecular weight standards (MW) are denoted and arrows indicate the specifically detected protein(s) for each antibody. Immunoblot experiments were repeated six times. **c**, 3-D SR-SIM of human cardiomyocytes containing RFP-labelled mitochondria (red) incubated with isolated GFP-labelled exogenous mitochondria (green) for 30 min. Coverslips were fixed and stained with a mitofusin-1 (MFN1) antibody (white). The top panels show the three separate color channels and the combined image (left to right). The boxed regions have been magnified in successive images (top right to lower panels, left to right) to demonstrate the reactivity of the MFN1 antibody with fusing mitochondria and the presence of this antibody throughout the cytosol. Parallel experiments were performed at 1, 2, and 4 h and the presented images are representative of four separate experiments. Scale bars, 0.5 μm.

Both cell types had equivalent amounts of MFN1 protein; however, only cardiomyocytes contained detectable quantities of MFN2. These findings suggest exogenous HCF mitochondria only have MFN1 on their outer membrane, which interacts with either MFN1 or MFN2 on endogenous cardiomyocyte mitochondria. Conversely, MFN2 did not appear to be necessary for fusion as exogenous HCF mitochondria efficiently fused with HCF endogenous mitochondria (Extended Data Fig. 3). Analysis of OPA1 expression revealed cardiomyocytes possess relatively small amounts of the long form of this protein as well as two short forms, while HCFs produced a larger quantity of only the long form. The long form of OPA1 is a membrane-anchored protein involved in fusion of the mitochondrial inner membrane, whereas the short forms are soluble proteins associated with mitochondrial fission and other cellular functions^28^. 3-D SR-SIM of MFN1 and MFN2 demonstrated that both of these proteins localised to fusing exogenous and endogenous mitochondria. At the same time, these mitofusin proteins were widely distributed throughout cardiomyocytes, which is consistent with their other known roles in the cell^26,27^. The presence of these proteins in cardiomyocyte nuclei may be indicative of a yet to be discovered function. Our results established that extracellular mitochondria are endocytosed in cardiac cells and most of these exogenous organelles end up fusing with the endogenous mitochondrial network (Extended Data Fig. 5). On the other hand, some exogenous mitochondria did not appear to escape from the endolysosomal compartments and are assumed to undergo hydrolysis and degradation through autophagy. This assumption is supported by the appearance of fragmented mitochondria in some lysosomes (Extended Data Fig. 2d).

## Discussion

The ability to distinguish the precise intracellular location of internalised exogenous human mitochondria using 3-D super-resolution and transmission electron microscopy in cardiac cells has allowed us to understand the intracellular trafficking of these organelles through time. Electron microscopy demonstrated that extracellular mitochondria begin to be endocytosed by human iPS-derived cardiomyocytes and primary cardiac fibroblasts within minutes. These organelles subsequently progress through the endolysosomal system from early endosomes to late endosomes to lysosomes. While some exogenous mitochondria appear to be destined for degradation in the latter compartment, our results show that the majority of exogenous mitochondria escape from endosomal and lysosomal compartments and then effectively fuse with endogenous cardiac cell mitochondria.

Our earlier experiments, and those of others, established that endocytosis of extracellular mitochondria was dependent on actin polymerization and that internalisation of these organelles resulted in long-term replenishment of mtDNA in HeLa cell-derived ρ^0^ cells, which are otherwise devoid of mtDNA^9,11,14,29–31^. Restoration of the ρ^0^ mitochondrial genome through uptake of mitochondria isolated from HeLa cells led to enhancement of intracellular ATP levels and oxygen consumption rates^9^. Because increases in ρ^0^ cellular respiration persisted for more than 50 population doublings, it was evident that endocytosed HeLa cell mitochondria containing functional mtDNA were lastingly integrated within these cells. However, we did not resolve how internalized HeLa cell mitochondria were transported within ρ^0^ cells and whether they fused with endogenous mitochondria, persisted as distinct organelles, or merely contributed their DNA to ρ^0^ mitochondria.

In this study, we defined the intracellular transit of extracellular mitochondria from the plasma membrane through to their incorporation with the endogenous mitochondrial network in two types of human heart cells — one dividing (HCFs) and one non-dividing (iPS-CMs). Our findings support the view that endocytosis and integration of exogenous mitochondria is an ancient biological process that is distinct from the recently described cell-to-cell transfer of mitochondria through tunneling nanotubes or exosomes^16,30,32,33^. In essence, our observations of the uptake of isolated mitochondria in human heart cells corroborates the endosymbiotic theory advanced and substantiated by Lynn Margulis (formerly Sagan) half a century ago^34^.

A complete understanding of the mechanisms that bring about endocytosis of extracellular mitochondria and fusion of those organelles with the endogenous mitochondrial network may provide a foundation for innovative new treatments to a number of human diseases and disorders^35^. Our earlier work has supported the use of mitochondrial transplantation to decrease infarction, increase ATP content, and improve functional performance of the ischemic heart^6–9,17^. We also demonstrated that delivery of exogenous mitochondria to the heart in animal models does not provoke an immune response nor produce arrhythmias^7,21^. Importantly, the clinical relevance and safety of autologous mitochondrial transplantation to the ischemic heart has recently been validated in five pediatric patients^36^. Aside from providing benefits to ischemic tissues, transplantation of exogenous mitochondria may also prove to be useful for treating mitochondrial myopathies and a number of other diseases with underlying mitochondrial dysfunction^13,15–18,20,37–40^. Further study of the molecular mechanisms of exogenous mitochondrial endocytosis and fusion with endogenous mitochondria are warranted and these inquiries may guide development of innovative treatments directed at augmenting impaired mitochondria in a wide range of tissues and cells.

## METHODS

### Reagents

The following antibodies were purchase from Abcam: anti-α-actinin (ab68167), anti-vimentin (ab92547), anti-TOMM20 (ab56783), anti-Mitochondria antibody [MTC02] (ab3298), anti-Mitochondria antibody [113-1] (ab92824), anti-Mitofusin 1 antibody (ab57602), anti-Mitofusin 2 antibody (ab101055), anti-OPA1 antibody (ab157457), anti-Rab5 antibody (ab18211), anti-Rab7 antibody (ab50533), and anti-Rab9 antibody (ab2810). The mouse IgG isotype control (10400C) and rabbit IgG isotype control (10500C) antibodies were purchased from ThermoFisher Scientific. The following CellLight reagents were purchased from ThermoFisher Scientific: Early endosomes (C10586 and C10587), Endoplasmic Reticulum (C10590 and C10591), Golgi Complex (C10592 and C10593), Late endosomes (C10588 and C10589), Lysosomes (C10596 and C10597), Mitochondria (C10600 and C10601) and Null Virus Negative Control (C10615). MitoTracker Red CMXRos (M7512) was purchased from ThermoFisher Scientific and used according to the manufacturer’s directions at 100 nM.

### Cells

iCell-Cardiomyocytes^2^ (iPS-CMs) (CMC-100-012-001) were purchased from Cellular Dynamics and Human Cardiac Fibroblasts (HCFs) (6300) were purchased from Sciencell Research Laboratories. Cells were cultured according to the manufacturer’s directions. Baculovirus infections were performed on dividing HCFs by adding 400 μL of CellLight reagent to a 100-mm confluent plate of cells for 16 h. Spontaneously contracting syncytial monolayers of iPS-CMs (≥ 96 hours after plating) were infected for 16 h by using 10% CellLight reagent in iCell Cardiomyocyte Maintenance Medium (CMM-100-120-001) (Cellular Dynamics) containing 0.2 μM BacMam Enhancer kit (ThermoFisher Scientific).

### Mitochondria

HCF media was replaced with 4 mL of 4°C homogenisation buffer (300 mM sucrose, 10 mM HEPES-KOH, 1 mM EGTA-KOH, pH 7.4) containing 2 mg Subtilisin A protease from *Bacillus licheniformis* (Sigma-Aldrich) and incubated at room temperature for 5 min. Digested cells were transferred to a 15-mL tube on ice and digested for an additional 15 min before filtration through a 10 μm mesh filter (PluriSelect) saturated with cold homogenisation buffer. Mitochondria were collected by centrifugation at 9,500 RCF at 4°C for 5 minutes and washed 3 times in cold homogenisation buffer before resuspension in culture media. Mitochondrial number was determined by using a Multisizer 4e Coulter Counter (Beckman-Coulter) and corroborated by hemocytometry. Isolated mitochondria were incubated with HCFs or iPS-CMs for 0 to 4 h.

### Labelling

Some isolated mitochondria (1 × 10^8^) were conjugated to 10 nm NHS-activated gold nanoparticles (CytoDiagnostics) according to the manufacturer’s directions. Labeled mitochondria were washed 5 times with homogenisation buffer containing 1 mg/mL fraction V bovine serum albumin (BSA) to remove unreacted nanoparticles. Gold-labeled mitochondria were incubated with iPS-CMs for 0, 2.5, 5 and 10 minutes prior to fixation and preparation for transmission electron microscopy.

### Immunoblots

Proteins were isolated by rinsing cells with ice-cold PBS (pH 7.4) and lifting them from the plates with a rubber policeman. Pelleted cells were resuspended in a small volume of ice-cold lysis solution (150 mM NaCl, 20 mM Tris-HCl (pH 7.6), 1 mM EDTA, 0.5% sodium deoxycholate, 70 mM NaF, 1% Nonidet P-40, Complete protease inhibitor cocktail (Sigma-Aldrich), 200 μM sodium orthovanadate, and 2 μM phenylmethylsulfonyl fluoride)^41^. After a 10-min incubation on ice with intermittent, brief agitation, debris was pelleted in a microcentrifuge, and the supernatants were stored at −80°C. Protein concentrations were determined using the bicinchoninic acid (BCA) protein determination kit (Pierce). SDS-PAGE and transfer to nitrocellulose was performed using the XCell SureLock Mini-Cell System and XCell II Blot Module (ThermoFisher), respectively. Identical gels were stained with Coomassie brilliant blue R250 to confirm equal protein loading. The ColorBurst Electrophoresis Marker (Sigma-Aldrich) and MagicMark XP Western Protein Standard (ThermoFisher) were used according to the manufacturer’s directions. Nitrocellulose membranes were rinsed in Tris-buffered saline (pH 7.4) containing 0.1% Tween 20 (TBS-T) and blocked in 5% non-fat milk in TBS-T for 1 h at 22°C on a rocking platform. Membranes were incubated overnight at 4°C on an orbital shaker with primary antibodies diluted 1:1000 in TBS-T containing 1% non-fat milk (Sigma). Antibodies were detected with the Amersham ECL Western Blotting Analysis System (GE Healthcare).

### Staining

Cells cultured on N° 1.5H (170 μm ± 5 μm) coverslips (Marienfeld-Superior) were fixed in 4% freshly-prepared formaldehyde in phosphate buffered saline (PBS). Cells were permeabilised in 0.1% Triton X-100 in PBS for 3 minutes and incubated with primary antibodies diluted 1:1000 in 10% fetal bovine serum (FBS) in PBS for 1 h. Primary antibodies were detected with species-appropriate Alexa Fluor 488-, 568-, or 633-conjugated secondary antibodies (ThermoFisher) diluted in PBS and cells were simultaneously stained using 4",6-diamidino-2-phenylindole (DAPI) (ThermoFisher). Coverslips were mounted to slides using ProLong Diamond Mountant (ThermoFisher).

### Microscopy

Cells were visualised on a ELYRA PS.1 (Zeiss) attached to a LSM 710 inverted microscope (Zeiss) with a 100x oil objective (Zeiss) and a DAPI/GFP/mRFP/Alexa 633 fluorescence filter set. Optical sections (84 nm) were collected through entire cell volumes using 5 grid patterns for structured illumination microscopy. Image stacks were processed using ZEN Black software (Zeiss) and displayed as transparent three-dimensional (3-D) volumetric renderings. Widefield fluorescence microscopy was performed using a FSX100 (Olympus). For transmission electron microscopy, iPS-CMs cultured on Thermanox coverslips (ThermoFisher) were fixed in 2% formaldehyde, 2.5% grade I glutaraldehyde, and 0.03% picric acid in 0.1 M cacodylate buffer, pH 7.4 at 4°C prior to incubation in 1% osmium tetroxide and 1.5% potassium ferrocyanide dissolved in cacodylate buffer. After contrast staining with 1% uranyl acetate in cacodylate buffer, cells were dehydrated and infiltrated with a 1:1 mixture of Epon-Araldite (Electron Microscopy Sciences) and propylene oxide. Embedding resins were polymerised for 24 to 48 h at 60°C and 60 nm-thick sections on were placed on copper grids and imaged on a Jeol 1200EX (80 kV).

### Biochemistry

ATP content was determined in the presence or absence of 1 μM adenosine diphosphate (ADP) using the ATPlite luminescence assay (PerkinElmer).

### Statistics

Control and experimental ATP data was expressed as concentration ± standard error of the mean (SEM). Statistical analysis was performed by Student’s t-test and a p < 0.05 was considered significant. The number of internalised GFP-labeled mitochondria associated with RFP-labeled early endosomes, late endosomes, lysosomes, or mitochondria was enumerated at 0.5, 1, 2, and 4 h by fluorescence microscopy. Twenty-five randomly-selected, high-powered (400x) fields were acquired at each time point and in each of the 4 cell compartments. From 1,852 to 12,007 mitochondria were counted per compartment at any given time. For early endosomes, late endosomes and lysosomes, GFP-labeled mitochondria were quantified as fully encapsulated, partially encapsulated (escaping) or not associated with each compartment. GFP-mitochondria were determined to be either fusing with the RFP-labeled endogenous mitochondria or not associated with these organelles. Analysis of the cell count data was performed using a generalised linear modeling approach with the gamma distribution and a log link function to evaluate the escaping mitochondria as a percentage of total cells between the 4 time points (0.5, 1, 2, and 4 h) within each compartment (early endosome, late endosome, and lysosome)^42^. Fit of the model was assessed by the Akaike information criterion. Summary data are expressed in terms of the mean percentage with the standard error and two-tailed Bonferroni adjusted. p < 0.05 are considered statistically significant based on the Wald test. Statistical analysis was conducted using SPSS Statistics version 23.0 (IBM).

### Data Availability

The datasets generated during and/or analysed during the current study are available from the corresponding author on reasonable request.

## Acknowledgements

Supported by the Boston Children’s Hospital Anesthesia Foundation, Ryan Family Endowment, Cardiac Conduction Fund, Richard and Susan Smith Foundation, President’s Innovation Award, Boston Children’s Hospital, Michael B. Klein and Family, Sidman Family Foundation, Kenneth C. Griffin Charitable Research Fund and Boston Investment Council. We also thank the Harvard Center for Biological Imaging for infrastructure and support and acknowledge NIH award 1S10RR029237-01, which was used to acquire the Zeiss ELYRA PS.1 microscope.

## Author contributions

D.B.C., P.J.D. and J.D.M. conceived of or designed the experiments. D.B.C., R.Y., and J.K.T. performed the experiments. D.B.C. R.Y., J.K.T. and D.Z. analysed the data. D.B.C. and J.D.M. wrote the manuscript. Correspondence and requests for materials should be addressed to D.B.C.

---

**Extended Data Figure 1.**
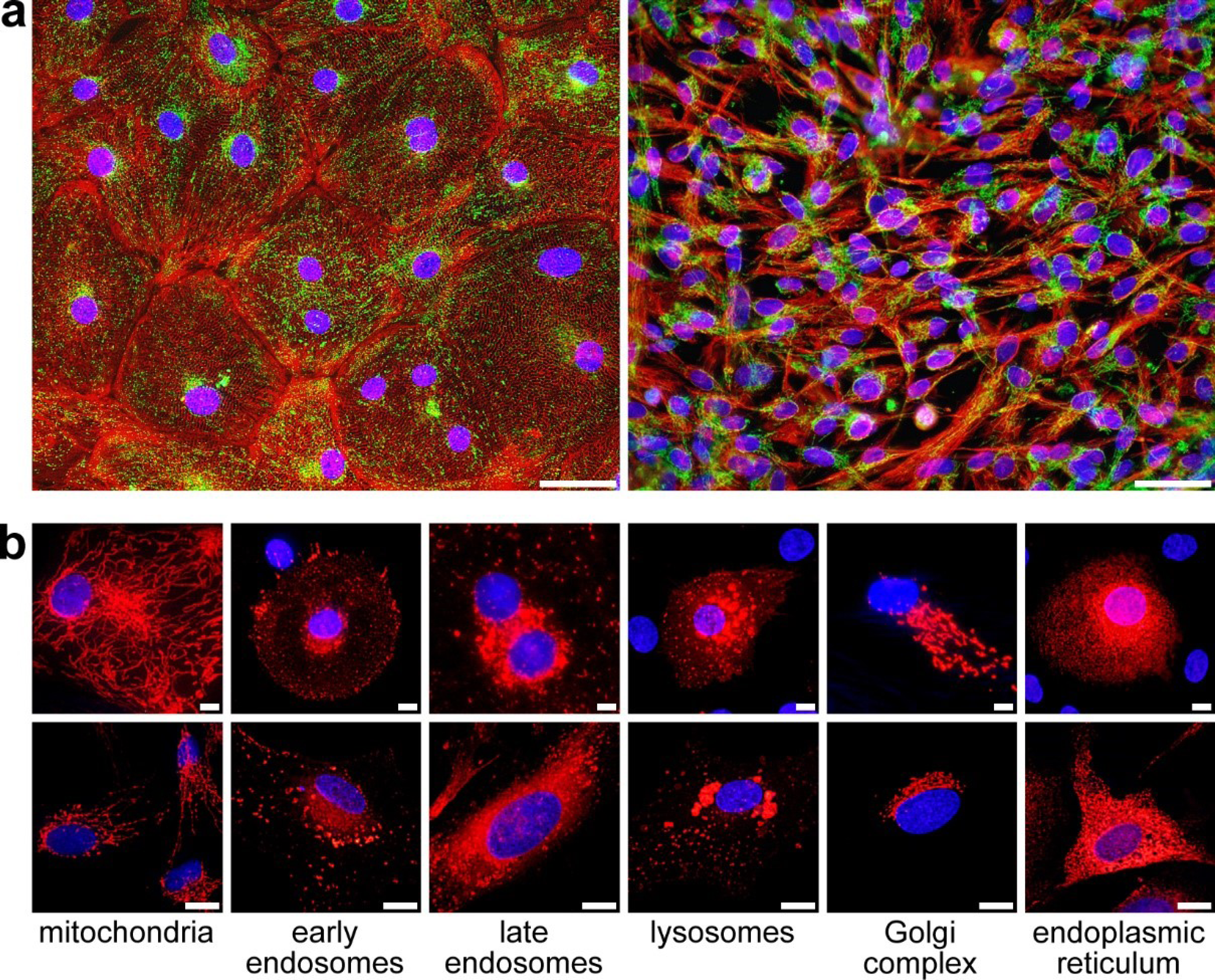
Widefield fluorescence microscopy of human cardiomyocytes and cardiac fibroblasts. **a**, iPS-CMs (left) and HCFs (right) stained with ACTN or vimentin antibodies (red), respectively. Mitochondria in these cells (green) were stained with MTC02 (left) or TOMM20 (right) and the nuclei were stained with DAPI (blue). Scale bars, 50 μm. **b**, Cardiomyocytes (top panels) and fibroblasts (bottom panels) infected with CellLight BacMam 2.0 reagents. Baculovirus infection rates after 16 h were similar for each cell type (84.45% ± 1.45 for iPS-CMs and 84.43% ± 1.80 for HCFs, mean percentage of cells expressing fluorescent proteins ± SEM). Expression of RFP was apparent in the appropriate cellular organelles. Images represent six separate infections with each baculovirus reagent. Scale bars, 10 μm.

**Extended Data Figure 2.**
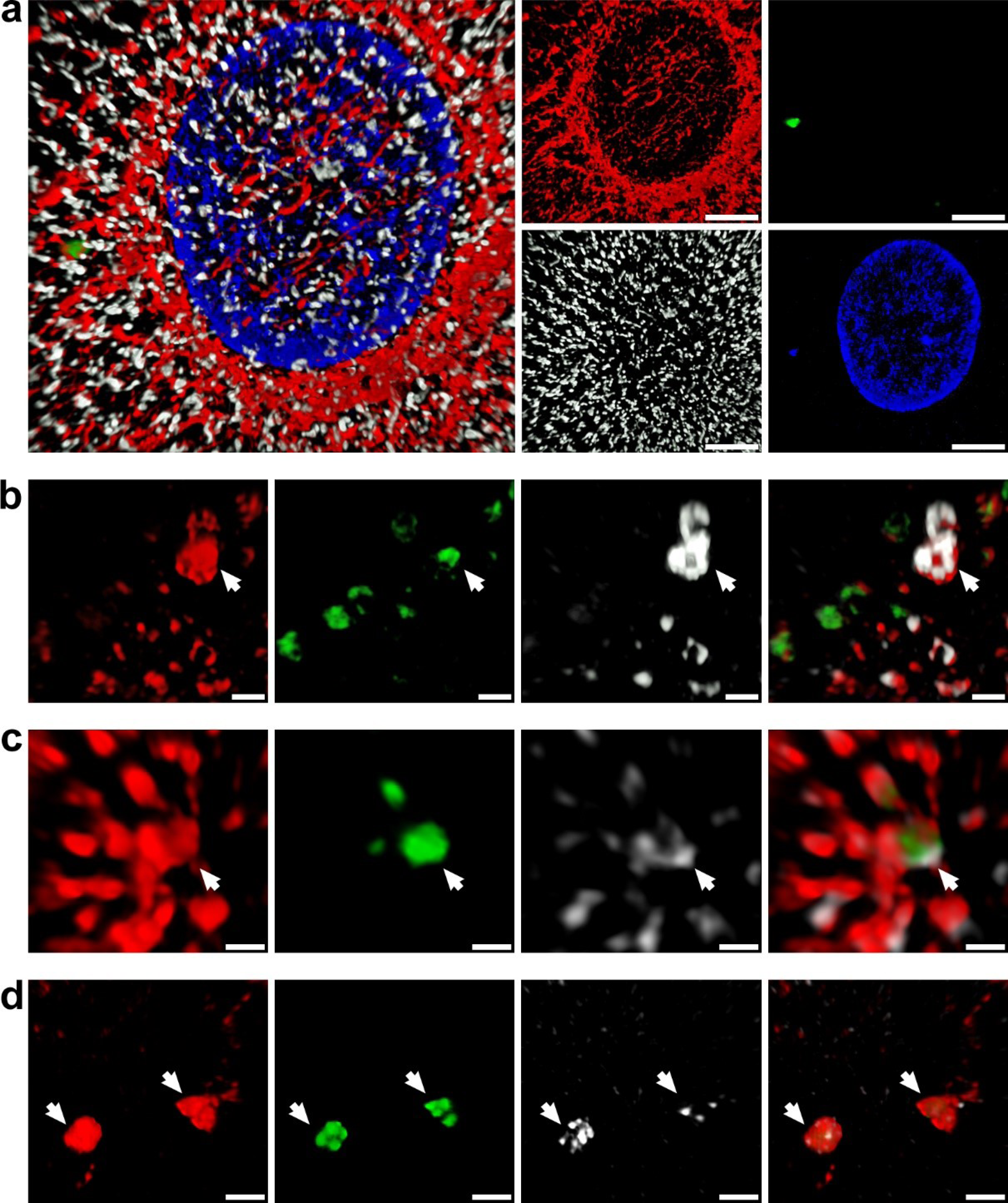
Internalised exogenous mitochondrial position within cardiomyocytes. **a**, An exogenous mitochondria (green) associated with the endogenous mitochondrial network (red) after 1 h in iPS-CMs that were subsequently stained for the contractile apparatus with ACTN (white) and DNA with DAPI (blue). The combined image (left) and individual colors (right) are shown. Images are representative of four separate experiments. Scale bar, 10 μm. **b**, Early endosomes (red), internalised mitochondria (green), and EEA1 staining (white) of iPS-CMs at 30 min. These images show exogenous mitochondria encapsulated within early endosomes, escaping from these vesicles, or not associated with this compartment. Images are representative of four separate experiments. Scale bars, 0.5 μm. **c**, Late endosomes (red), internalised mitochondria (green), and Rab7 staining (white) of iPS-CMs at 2 h. These panels show exogenous mitochondria encapsulated and escaping from late endosomes. Images are representative of four separate experiments. Scale bars, 0.5 μm. **d**, Lysosomes (red), internalised mitochondria (green), and Rab7 staining (white) of iPS-CMs at 2 h. The encapsulated GFP-labelled mitochondria appear to be fragmented, indicative of hydrolytic enzyme activity. Images are representative of four separate experiments. Scale bars, 0.5 μm.

**Extended Data Figure 3.**
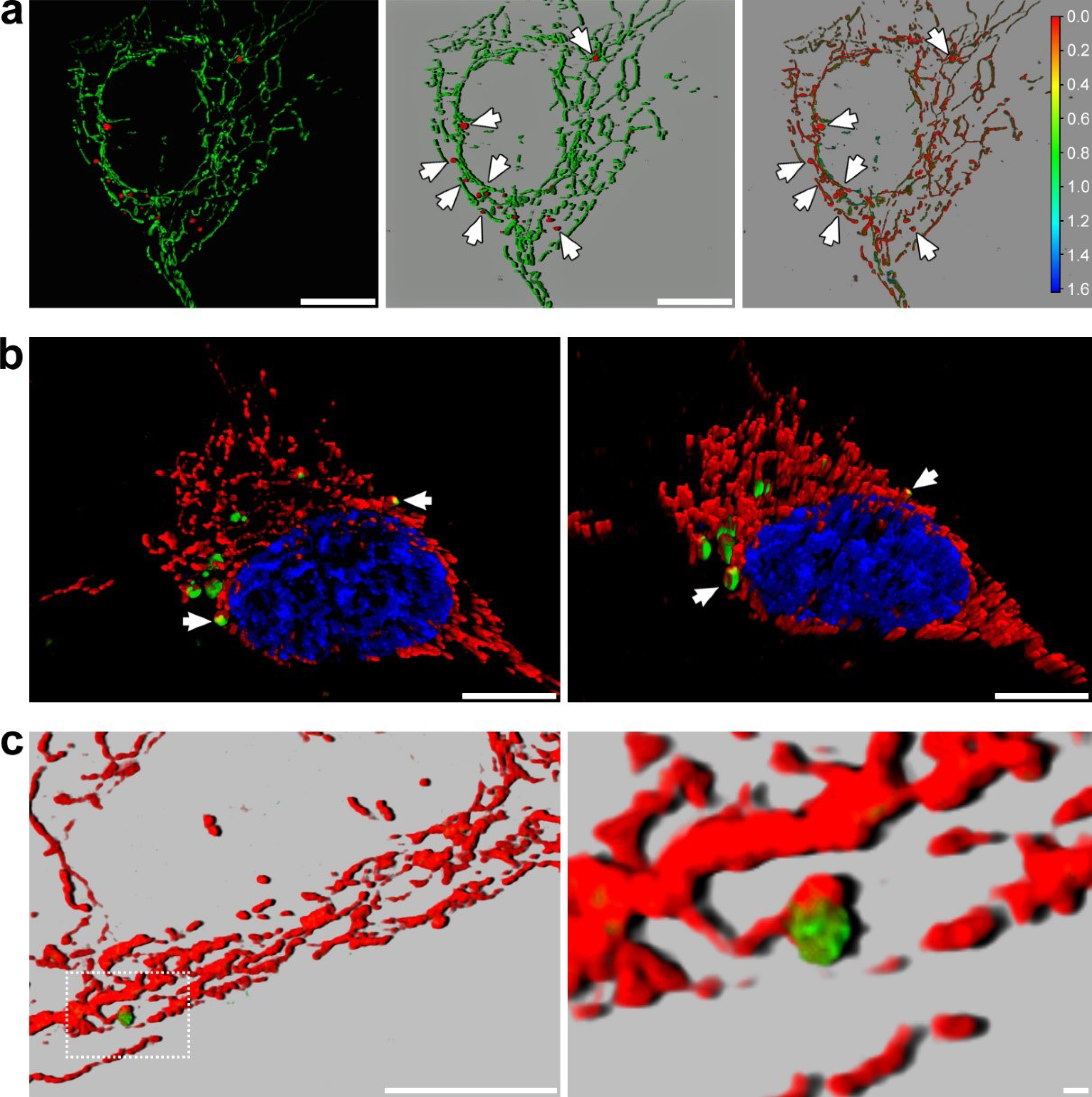
Exogenous mitochondrial fusion with endogenous mitochondria in human cardiac fibroblasts. **a**, RFP-labelled HCF mitochondria were isolated and incubated with HCFs containing GFP-labelled mitochondria for 2 h. The transparent rendering was depicted with a black (left panel) and grey background (middle panel) to accentuate the association of exogenous mitochondria with the endogenous mitochondrial network (arrows). Depth coding analysis using Zen Black software (Zeiss) showed that RFP and GFP mitochondria occupied the same spatial position in the 1.6 μm thick cell. The color scale indicates distance (μm) from the bottom to the top of the cell. Images are representative of four separate experiments. Scale bars, 10 μm. **b**, a 3-D volumetric rendering (left) and rotation (right) of a fibroblast containing RFP-labelled mitochondria that had been exposed to isolated GFP-labelled HCF extracellular mitochondria for 4 h. Fusion of exogenous and endogenous mitochondria was apparent (arrows). Images are representative of four separate experiments and similar results were obtained at 0.5, 1, and 2 h. Scale bars, 10 μm. **c**, fusion of an exogenous GFP-labelled mitochondria with the HCF mitochondrial network (red) at 1 h. 3-D volumetric renderings are depicted with a grey background to emphasize the colors. The boxed region in the left image is magnified on the right. Scale bars, 0.5 μm.

**Extended Data Figure 4.**
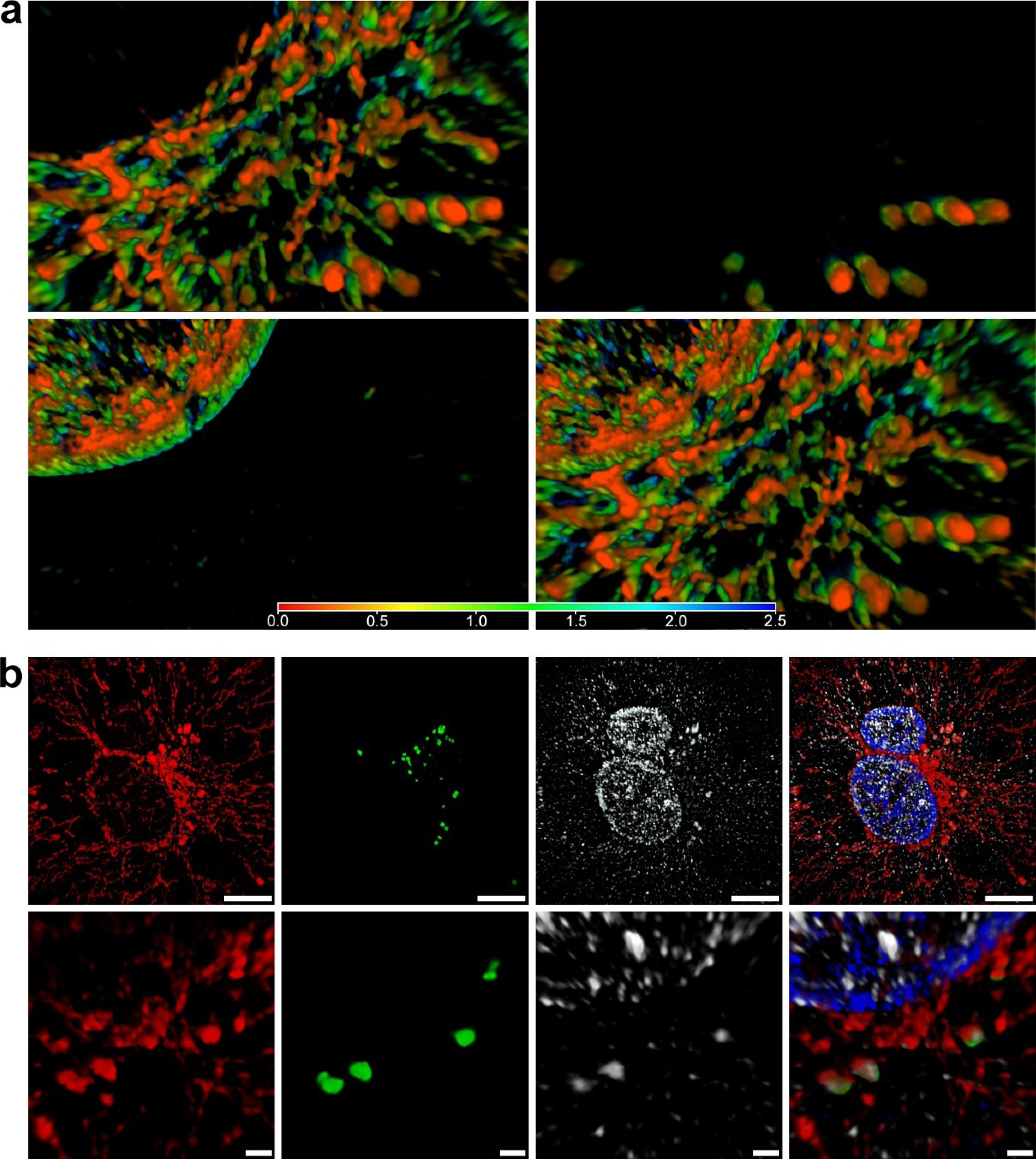
Depth-coding analysis of fusion in iPS-CMs and involvement of mitofusin-2. **a**, Depth coding analysis of the images presented in Fig 4a using Zen Black software (Zeiss). The color scale extends for 2.5 μm and the fusing mitochondria occupied identical spatial positions. **b**, 3-D SR-SIM of human cardiomyocytes containing RFP-labelled mitochondria (red) incubated with isolated GFP-labelled exogenous mitochondria (green) for 2 h. Coverslips were fixed and stained with a mitofusin-2 (MFN2) antibody (white). The top and bottom panels show separate color channels and the combined image at two different magnifications. Although the antibody reacted with fusing mitochondria, there was also considerable staining of the nucleus and cytosol that was not associated with mitochondria. Images are representative of four separate experiments. Scale bars, 10 μm (top) and 0.5 μm (bottom).

**Extended Data Figure 5.**
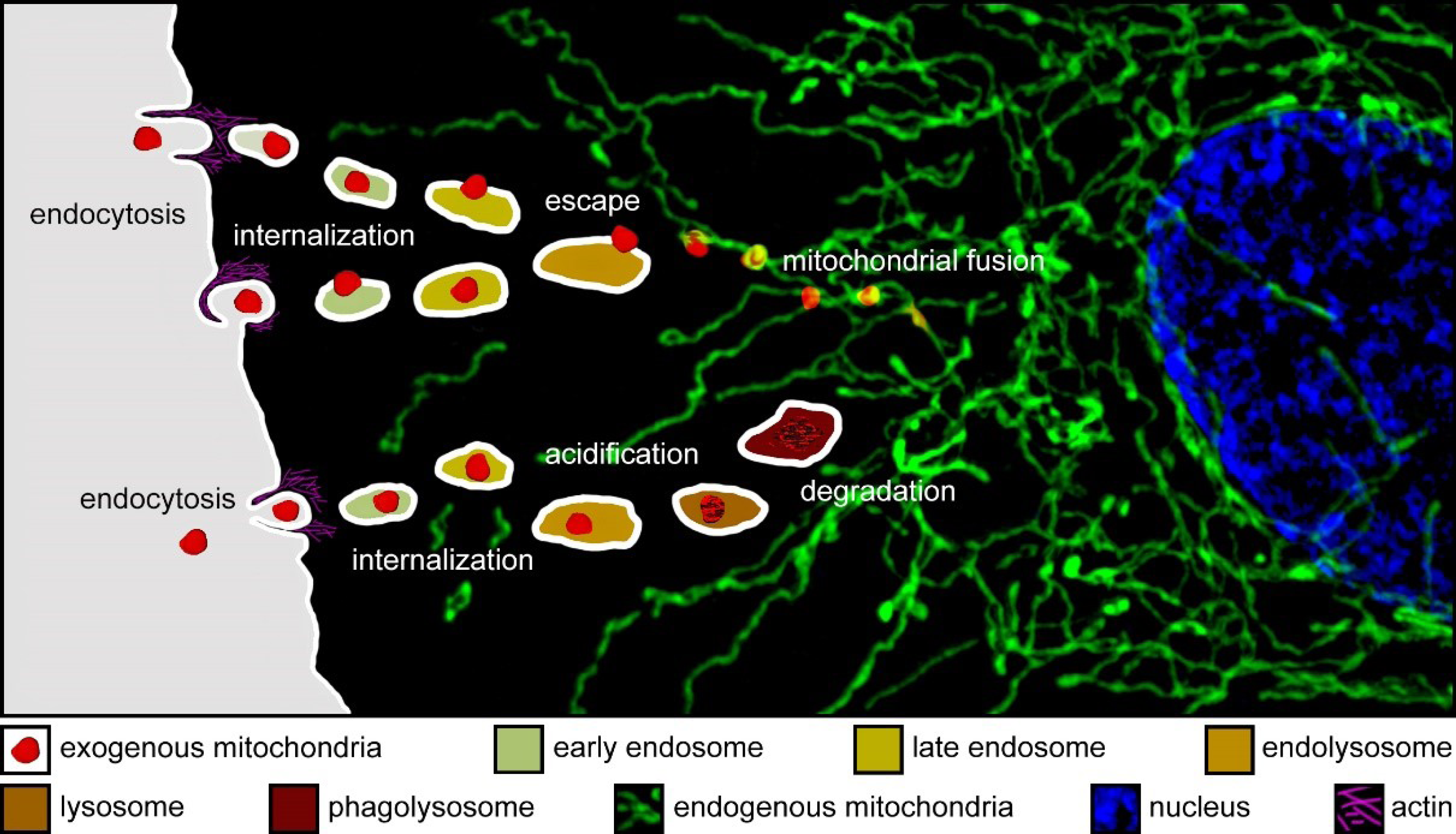
Schematic representation of the fate of endocytosed mitochondria. Extracellular mitochondria are internalised in human cardiomyocytes and cardiac fibroblasts through actin-dependent endocytosis. These exogenous mitochondria escape from endosomal and lysosomal compartments and fuse with the endogenous mitochondrial network or are degraded through hydrolysis and phagocytosis.

